# T6SS4 is heterogeneously expressed in *Y. pseudotuberculosis* and is a target for transcriptional and post-transcriptional regulation

**DOI:** 10.1101/2025.07.10.664076

**Authors:** Anna Kerwien, Britta Körner, Ines Meyer, Yannick Teschke, Cassandra S. Köster, Ileana P. Salto, Petra Dersch, Anne-Sophie Herbrüggen

**Affiliations:** Institute for Infectiology, Center for Molecular Biology of Inflammation (ZMBE), University of Münster, Münster, Germany

## Abstract

The type VI secretion system (T6SS) is a complex secretion system encoded by many Gram-negative bacteria to translocate effector proteins directly into target cells. Due to its high complexity and energy-intensive firing process, regulation of the T6SS is tightly controlled in many organisms. *Y. pseudotuberculosis* encodes four complete T6SS clusters but lacks genes implicated in T6SS gene regulation in other microorganisms, indicating a distinct control mechanism. Here, we could show that the T6SS4 of *Y. pseudotuberculosis* is heterogeneously expressed within a population, which is determined by the transcriptional T6SS4 activator RovC. Moreover, the T6SS4 and RovC are embedded in a complex and global regulatory network, including the global post-transcriptional regulator CsrA, the *Yersinia* modulator A (YmoA), the global protease Lon, and RNases (PNP and RNase III). Post-transcriptional processing of the T6SS4 polycistron and different transcript stability within the operon also achieve a higher regulatory complexity. In summary, our work provides new insights into the sophisticated and complex regulatory network of the T6SS4 of *Y. pseudotuberculosis*, which clearly differs from regulation in other organisms.

**Authors summary:** Bacteria use a specialized multi-protein complex called the Type VI secretion system (T6SS) to inject toxic proteins into other cells to compete with target microorganisms or to infect host organisms. While the T6SS has been extensively studied in some model organisms, much less is known about the function and regulation of the four T6SS clusters of the food-borne human pathogen *Yersinia pseudotuberculosis*. In this study, we found that the T6SS4 of *Y. pseudotuberculosis* is only expressed in a small subpopulation *in vitro*. This suggests that its regulation is fundamentally different from what is known in other organisms. We show that a complex regulatory network regulates T6SS4 gene expression, and the T6SS4 transcript is post-transcriptionally processed, resulting in different mRNA levels of the individual T6SS components. These findings contribute to a deeper understanding of how bacteria, especially *Y. pseudotuberculosis,* regulate complex secretion systems at multiple levels.

## Introduction

Bacteria in complex environmental settings are exposed to various rapidly changing conditions, such as temperature and nutrient availability, competing with other microorganisms or evading the mammalian immune system. *Yersinia pseudotuberculosis* can be found in soil, water, plants and is a human pathogen that causes various gut-associated symptoms such as diarrhea and enteritis [1–4]. Entry into the human host involves a significant temperature and nutrient composition change. Moreover, the bacteria have to switch from defending a niche against other microorganisms to evading the host’s immune system, which requires a rapid and precise change in gene expression of the respective virulence factors. Due to these distinct lifestyles, it is not surprising that the expression of virulence-associated and fitness-relevant genes of *Y. pseudotuberculosis*, including its Type III (T3SS) and Type VI secretion systems (T6SS), is tightly regulated by temperature [5,6]. One well-described example is the regulation of the *Y. pseudotuberculosis* Ysc-Type III secretion system (Ysc-T3SS). The Ysc-T3SS promotes the injection of antiphagocytic Yop (*Yersinia* outer protein) effector proteins into host cells and plays an important role in defending against the attack of phagocytic cells [7–11]. Expression of the Ysc-T3SS is activated by the transcriptional regulator LcrF (low <u>c</u>alcium-response <u>r</u>egulator) at 37°C via an RNA-thermometer and is transcriptionally repressed at 25°C by the *<u>Y</u>ersinia* <u>mo</u>dulator <u>A</u> (YmoA) [12–17]. YmoA acts as a protein thermometer as it undergoes a conformational change and is rapidly degraded by the Lon/ClpP proteases upon a temperature upshift from moderate (25°C) to body temperature (37°C) [14,16].

In contrast to the regulation of the T3SS, much less is known about the regulation of the four T6SS clusters of *Y. pseudotuberculosis,* differing in their chromosomal arrangement and number of genes [18]. Exclusively encoded in Gram-negative bacteria, T6SSs are large contact-dependent secretion systems of 13 highly conserved core components that directly translocate a broad spectrum of different effector proteins into competing prokaryotic or eukaryotic cells [19–25]. As the assembly and disassembly of the apparatus are very energy-consuming, we expected that the expression of T6SSs of *Y. pseudotuberculosis* would be tightly regulated in response to environmental signals, as observed in other microorganisms [26–29]. Previous studies on the T6SS4 gene cluster of *Y. pseudotuberculosis* revealed that its expression is strongly temperature-dependent. It can only be induced at moderate growth temperature (25°C), predominantly during the stationary phase, but not at 37°C *in vitro* [18,30]. It was also found that expression of T6SS4 is promoted by direct binding of global regulators such as RpoS, OmpR, or RovM, and T6SS4 expression is positively regulated by quorum sensing [18,31–34]. Unlike any other yet known T6SS, *Yersinia*-T6SS4 gene expression further requires the expression of the specific hexameric transcriptional activator RovC [30]. RovC is encoded in the opposite direction upstream of the T6SS4 cluster and activates T6SS4 gene expression by direct binding within the T6SS4 promoter region [30]. The T6SS4 of *Y. pseudotuberculosis* is additionally controlled by the global regulator CsrA (<u>c</u>arbon <u>s</u>torage <u>r</u>egulator <u>A</u>) [30]. The CsrA protein belongs to the Csr system, which includes two small non-coding RNAs CsrB and CsrC. They can bind and sequester multiple CsrA proteins, thus preventing CsrA from binding to its target mRNAs. The Csr system is known to be an important post-transcriptional regulatory system that influences the expression of many virulence- and fitness-relevant genes in bacteria [35–38]. In *Y. pseudotuberculosis*, CsrA was found to play an essential role in T6SS4 expression, as it affects *rovC* on the transcriptional and post-transcriptional levels [30]. Although the exact mechanism of how CsrA influences RovC synthesis is still unclear, CsrA was shown to repress the *rovC* transcription indirectly, but also to stabilize *rovC* mRNA, indicating a complex role in the regulation of RovC and, thus, of T6SS4 [30]. It is further known that the CsrA homologous protein RsmA (<u>r</u>egulator of <u>s</u>econdary <u>m</u>etabolites <u>A</u>) negatively regulates the expression of all three T6SS islands in *Pseudomonas aeruginosa*, highlights the critical role of the Rsm/Csr system in T6SS regulation [39].

As several studies have shown that the expression of the T6SS4 is repressed at 37°C [18,30,31], it can be assumed that T6SS4 targets (micro-)organisms other than mammals, *e.g.*, competing bacteria in their environmental niches. However, no antibacterial effector that is exclusively translocated by T6SS4 has yet been identified. In contrast, it was hypothesized that *Y. pseudotuberculosis* uses its T6SS4 to maintain the intracellular ion homeostasis of *e.g.*, manganese or zinc [40–43]. This suggests a role in the resistance to oxidative stress and could promote a higher virulence in mice [40,42,43]. However, no expression of the T6SS4 was detectable by RNA-sequencing under different virulence-relevant conditions at 37°C and during infection in other studies [6,44–47].

To gain further insight into the role of the T6SS4, its temperature control mechanisms, and function, we analyzed the expression control of the *Y. pseudotuberculosis* T6SS4 at transcriptional, post-transcriptional, and translational levels. To this end, we used a flow cytometry-based approach to study T6SS4 expression at a single-cell level. We found that T6SS4 cluster expression within a population is very heterogeneous, unlike T6SS gene clusters in *Vibrio* or *Pseudomonas* spp [19,22,23,26,48–50]. We could further show that the heterogeneous expression of the transcriptional regulator gene *rovC* promotes phenotypic heterogeneity. Moreover, heterogeneous *rovC* and T6SS4 gene expression are impacted by the global protease Lon, the RNases PNPase and RNase III, the RNA-binding protein CsrA, and the transcriptional regulator YmoA. The rapid downregulation of *rovC* and T6SS4 gene expression upon temperature shift from 25°C to 37°C involves distinct temperature-dependent post-transcriptional modifications of both *rovC* and T6SS4 mRNA.

## Results

### Flow cytometry-based method revealed heterogeneous expression of the T6SS4 at different temperatures

In previous studies, T6SS4 expression of *Y. pseudotuberculosis* was mainly studied in bulk approaches, revealing only an overall up- or downregulation of gene expression in a bacterial population [18,31,34]. In contrast to this general approach to analyze gene expression, we applied flow cytometry and fluorescence microscopy to study T6SS4 expression at a single-cell level. Therefore, we used a strain in which the core gene *clpV4* of the T6SS gene cluster 4 (T6SS4) is chromosomally fused to *gfp* (**Fig 1A**). We could show that only a small subpopulation (approximately 10-15%) highly expressed *clpV4-gfp* (T6SS4-ON subpopulation), whereas the majority of the population remained repressed (T6SS4-OFF subpopulation) (**Fig 1B, S 1 Fig.**). This finding of phenotypic heterogeneity—referring to variable T6SS4 expression within a genetically identical population—differs from what was found in *P. aeruginosa* and *Vibrio cholerae*. In these model organisms, T6SS gene cluster expression was reported to be homogeneous among genetically identical cells [22,48,51–53]. To analyze a higher bacterial cell number, a flow cytometry-based method was established by gating for GFP-positive (GFP^+^, T6SS4-ON) and negative (GFP^-^, T6SS4-OFF) subpopulations (**Fig 1C, S 2 Fig**). Incubation at 25°C resulted in an increasing amount of *clpV4-gfp* expressing bacteria after 8 h, when the cultures reached the stationary phase. In contrast, *clpV4-gfp* expression was rapidly downregulated after a temperature shift to 37°C and remained repressed throughout the bacterial growth phase (**Fig 1B, C**). This aligns with previous findings, describing a strong regulatory effect of temperature and growth phase on the T6SS4 expression [18]. It also illustrates that the overall induction of the T6SS4 expression is only triggered in a subset of the population.

**Fig 1.**
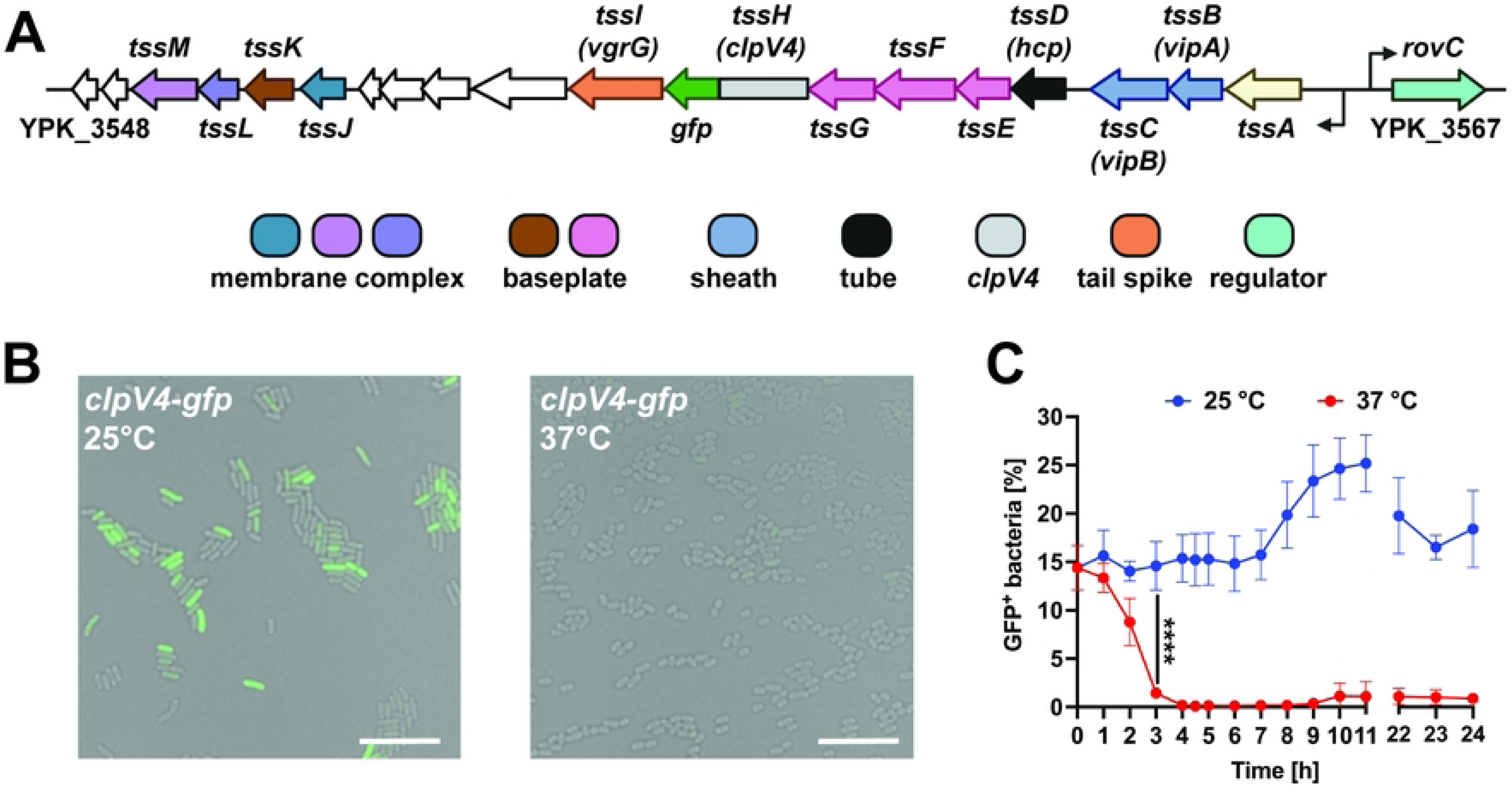
T6SS4 of *Y. pseudotuberculosis* YPIII is heterogeneously expressed. (A) Scheme of YPIII T6SS4 cluster. Chromosomal fusion of *clpV4* to *gfp* was used as a reporter to analyze expression of T6SS4 genes. (B) Fluorescence microscopy of wt *clpV4-gfp*. The bacteria were incubated for 6 h at 25°C or 37°C and imaged on 1% agarose pads. The scale bar represents 10 μm, and representative overlays of the GFP channel and brightfield are shown. (C) Quantification of *clpV4-gfp* expressing bacteria (GFP^+^) incubated at 25°C or 37°C. 1 x 10^5^ bacteria were analyzed using flow cytometry. Data depict the mean and standard deviation of three independent experiments. Statistical differences were determined using a Two-Way ANOVA with Šidák correction. **** = p ≤ 0.0001.

### The heterogeneous expression of the transcriptional regulator gene *rovC* causes heterogeneity of T6SS4

Our previous study revealed that expression of T6SS4 genes is activated by the hexameric transcriptional regulator RovC. It was further shown that deletion of *rovC* completely abolishes T6SS4 expression at 25°C [30]. Therefore, we assumed that *rovC* might only be expressed in the T6SS4-ON subpopulation. To test this hypothesis, we introduced a low-copy-number plasmid containing the translationally fused promoter region of *rovC* to *mCherry* (**S 2 Fig.**) into the *Y. pseudotuberculosis* strains expressing the chromosomally encoded *clpV4-gfp* fusion. With this dual reporter strain, the expression of *rovC-mCherry* and *clpV4-gfp* was analyzed simultaneously. As shown in **Fig 2A**, the *rovC-mCherry* fusion was weakly expressed in the majority of the bacteria (mCherry^low^ population), and these bacteria did not express *clpV4-gfp* (**Fig 2B, C**). In contrast, 15-20% of the bacteria showed a high expression level of *rovC-mCherry* and expressed *clpV4-gfp*. This strongly indicates that the heterogeneous expression of the transcriptional regulator gene *rovC* causes heterogeneity of T6SS4 expression.

**Fig 2.**
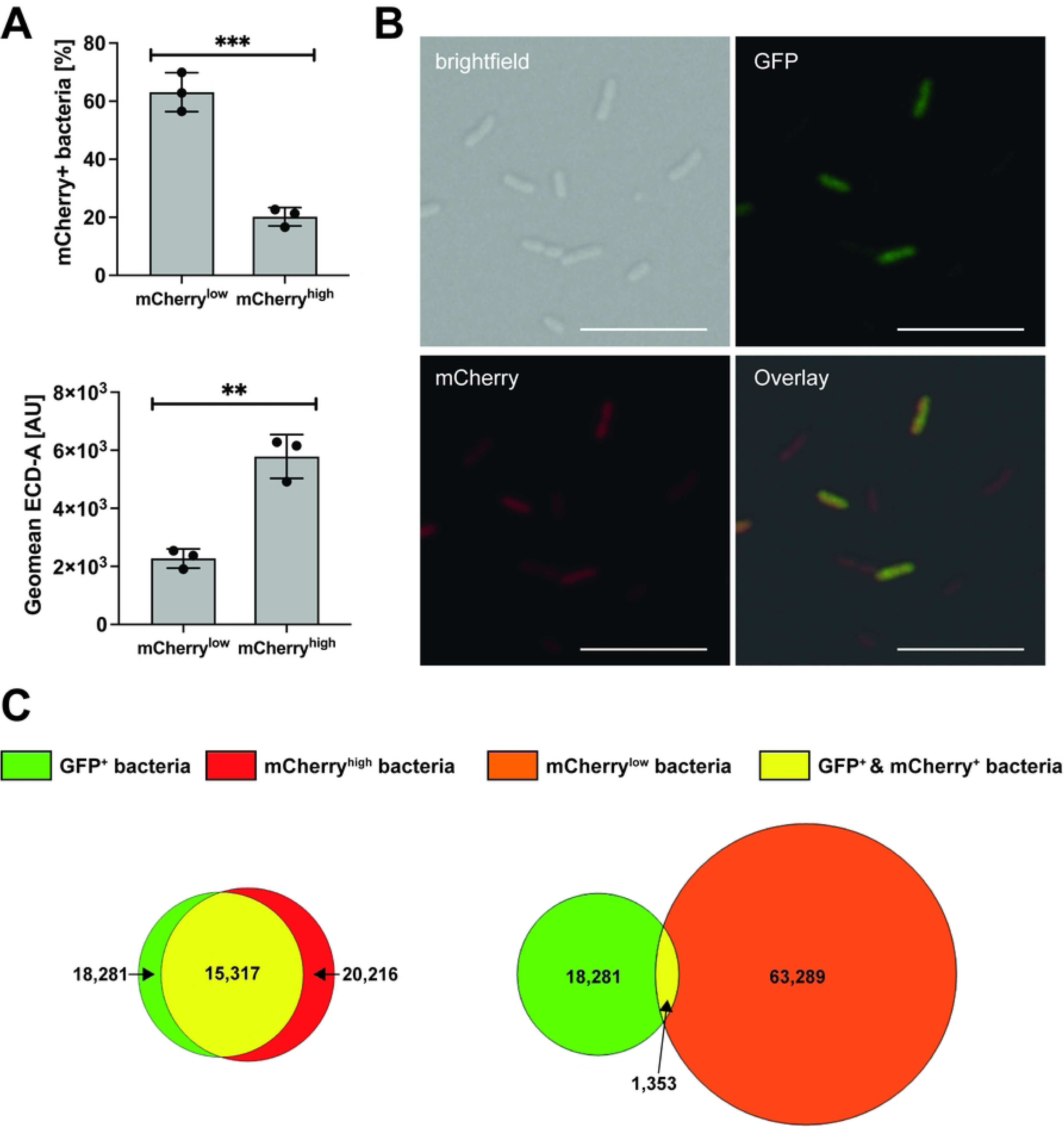
Heterogeneous T6SS4 expression is determined by RovC. (A) Flow cytometric analysis of a translational *rovC* promoter fusion to mCherry (p*P_rovC_ rovC’-‘mCherry*) resulted in two subpopulations of mCherry^low^ and mCherry^high^ expression, which differed in their expression intensity (GeoMean ECD-A). Significant differences were determined using an unpaired t-test. (B) Fluorescence microscopy of *Y. pseudotuberculosis* YPIII *clpV4-gfp* pP*_rovC_ rovC’-‘mCherry*. Images of brightfield, GFP, and mCherry channel are shown, as well as an overlay of all channels. Representative images from one of three independent experiments are shown. Scale bar indicates 10 μm. (C) Quantitative Venn diagram of *mCherry* and *gfp* expressing bacteria, analysed by flow cytometry of wt *clpV4-gfp* with translational fusion of promoter region of *rovC* to *mCherry* (pP*_rovC_ rovC’*-‘*mCherry*). 1 x 10^5^ bacteria were analysed. Data depict the mean of three independent experiments. Absolute numbers of GFP-expressing bacteria (green), mCherry^high^-expressing bacteria (red), mCherry^low^-expressing bacteria (orange), and bacteria expressing both (yellow). ** = p ≤ 0.01, *** = p ≤ 0.001.

### Influence of the global post-translational, post-transcriptional, and transcriptional regulators Lon, CsrA, and YmoA on temperature-dependent *rovC* and T6SS4 gene expression

Next, we aimed to gain insight into the molecular mechanisms of how heterogenous T6SS4 expression is controlled in response to temperature, *e.g.*, which factors contribute to the rapid decrease of the number of T6SS4-ON bacteria in the population upon a temperature shift from 25°C to 37°C. Several global regulators of *Yersinia* have been shown to control gene expression in a temperature-dependent manner. One example is the global protease Lon [54–59], which was shown to be involved in the temperature-dependent degradation of the virulence regulators YmoA and RovA of *Y. pestis* and *Y. pseudotuberculosis,* respectively [60,61]. We first examined whether the Lon protease affects the number of T6SS4-ON bacteria in a temperature-dependent manner. We found that deleting the *lon* gene substantially increased *clpV4-gfp* (T6SS4-ON)-expressing bacteria at 25°C compared to the wildtype (**Fig 3A**). However, the number of the *clpV4-gfp* (T6SS4-ON)-expressing bacteria decreased rapidly when the culture was shifted from 25°C to 37°C. Based on this result, it is possible that Lon directly targets RovC or influences *rovC* transcription indirectly (**Fig 3B**). Western blot analysis demonstrated that the levels of T6SS4 components, such as Hcp4 and ClpV4, as well as the T6SS4 activator RovC are strongly increased in the absence of Lon at 25°C but not at 37°C (**Fig 3C-D**). This indicates that Lon exerts its influence on T6SS4 gene expression via the regulation of RovC. We further tested the impact of the *lon* gene deletion on the number of *mCherry^high^* (RovC-ON) bacteria in *Y. pseudotuberculosis* expressing the *P_rovC_rovC’-‘mCherry* reporter (**S 3 Fig**.) and the overall amount of the *rovC* transcript in the bacterial cells (**Fig 3E**). We found a higher number of RovC-ON bacteria and a significant increase in *rovC* transcript levels in the *lon* deletion mutant at 25°C. In contrast, the *rovC* transcript was downregulated at 37°C independently of the presence of Lon (**Fig 3E**), indicating that Lon controls the overall level of RovC but is not involved in temperature control.

**Fig 3.**
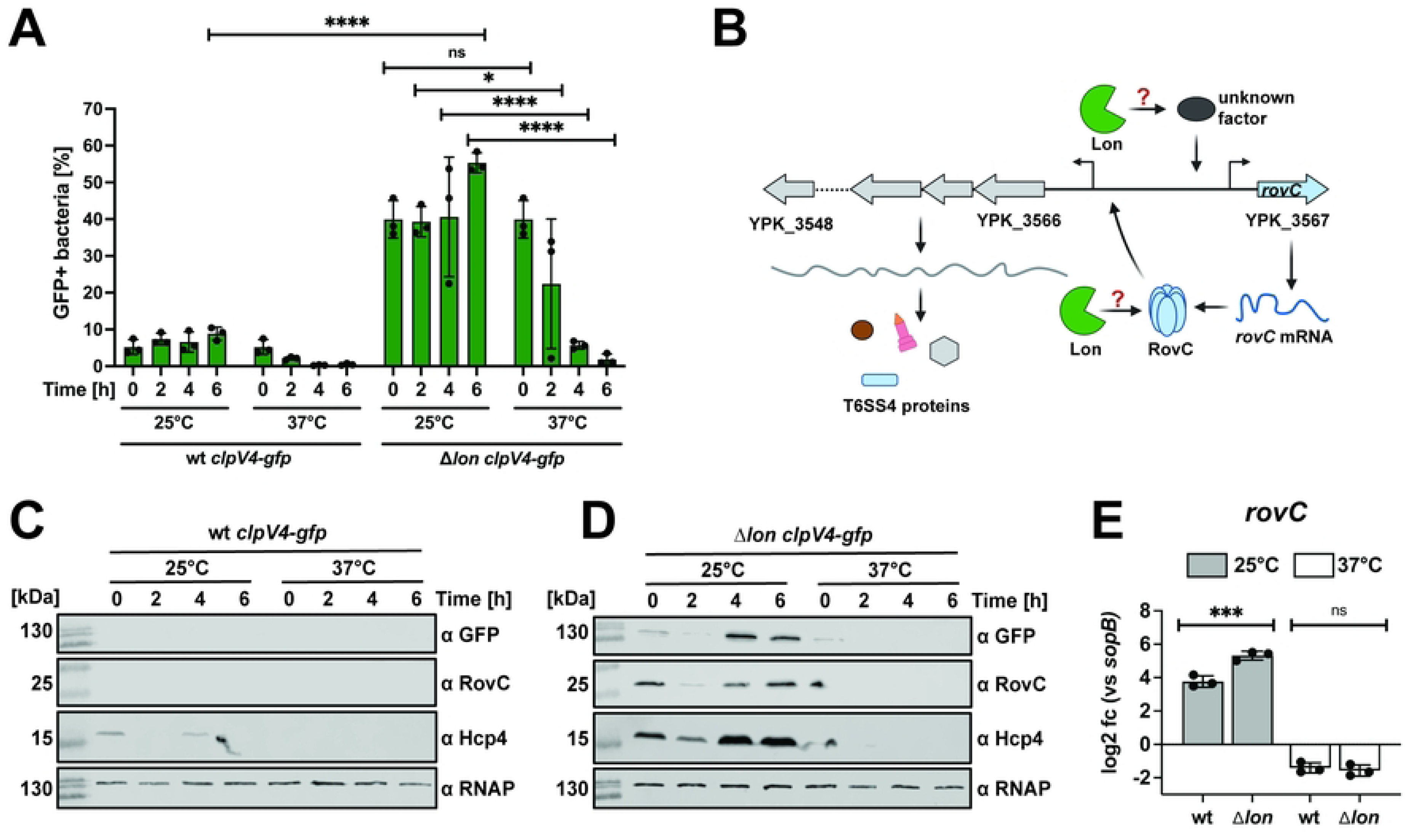
Temperature-based downregulation is not due to Lon-mediated degradation at 37°C. (A) Quantification of *clpV4-gfp* expressing bacteria in wt *clpV4-gfp* (YP412) and Δ*lon clpV4-gfp* (YP544) at either 25°C or 37°C. 1 x 10^5^ bacteria were analyzed using flow cytometry. Data depict the mean and standard deviation of three independent experiments. Significant differences were determined using a Two-Way ANOVA with Šidák correction. (B) Scheme of potential downregulation at 37°C mediated by Lon-dependent inhibition of the *rovC* transcription or by Lon-mediated proteolysis of RovC. Created in BioRender. Dersch, P. (2025) https://BioRender.com/8el9map. (C-D) Western blotting of wt *clpV4-gfp* and Δ*lon clpV4-gfp*. Protein levels of ClpV4-GFP, RovC, and Hcp4 were detected using antibodies against GFP, RovC, and Hcp4. Detection of RNAP was used as a loading control. Experiments were performed in three independent replicates; one representative Western blot is shown. (E) qRT-PCR was performed of total RNA extracted from wt and Δ*lon* incubated for 6 h at 25°C or 37°C. Specific primer pairs were used to determine expression levels of *rovC,* and log2 fold changes to *sopB* as a non-temperature-regulated reference gene [6] were calculated. Experiments were performed in three independent replicates, and the mean and standard deviations are shown. Significant differences were determined using an unpaired t-test. * = p ≤0.05, *** = p ≤ 0.001, **** = p ≤ 0.0001, ns = not significant p > 0.05.

Previous work has shown that the expression of the Csr system components of *Yersinia,* which are known to repress RovC and thus T6SS4 synthesis [30], is strongly controlled by temperature and carbon/nutrient sources [16]. CsrA regulates RovC synthesis in opposing ways: it represses transcription while promoting post-transcriptional stability [30]. Moreover, a *Yersinia*-specific histone-like protein YmoA, with homology to the *E. coli* Hha protein, is known to regulate *Yersinia* virulence factors, including the master regulator LcrF of the Ysc-T3SS gene cluster in a temperature-dependent manner [12,13]. From a transcriptomic analysis, it is further known that YmoA also influences the expression of several virulence-associated genes, such as *vipA4*, *vipB4,* and *rovC* in *Y. pseudotuberculosis*, and modulates the expression of the Csr system [12,15,16]. Based on this knowledge, we used *csrA* and *ymoA* deletion strains to test their influence on the synthesis of RovC and T6SS4 components at 25°C and 37°C (**Fig 4A**). A deletion of *csrA* or *ymoA* resulted in a significant increase in the number of T6SS4-ON bacteria at 25°C (**Fig 4B**), whereby the overall expression was significantly higher in the Δ*csrA* compared to the Δ*ymoA* mutant strain (**Fig 4B, C**). In addition, both gene deletions also led to a substantial increase in the RovC and Hcp4 levels at 25°C compared to wt (**Fig 4D**). However, no upregulation of RovC and the T6SS4 components was observed in the mutant strains at 37°C (**Fig 4B-D**). A qRT-PCR analysis further showed that *rovC* and T6SS4 gene transcript levels decreased significantly between 25°C and 37°C in both mutants (**Fig 4E**, **S 4 Fig.**), indicating that CsrA and YmoA affect RovC and T6SS4 expression but are not involved in temperature control.

**Fig 4.**
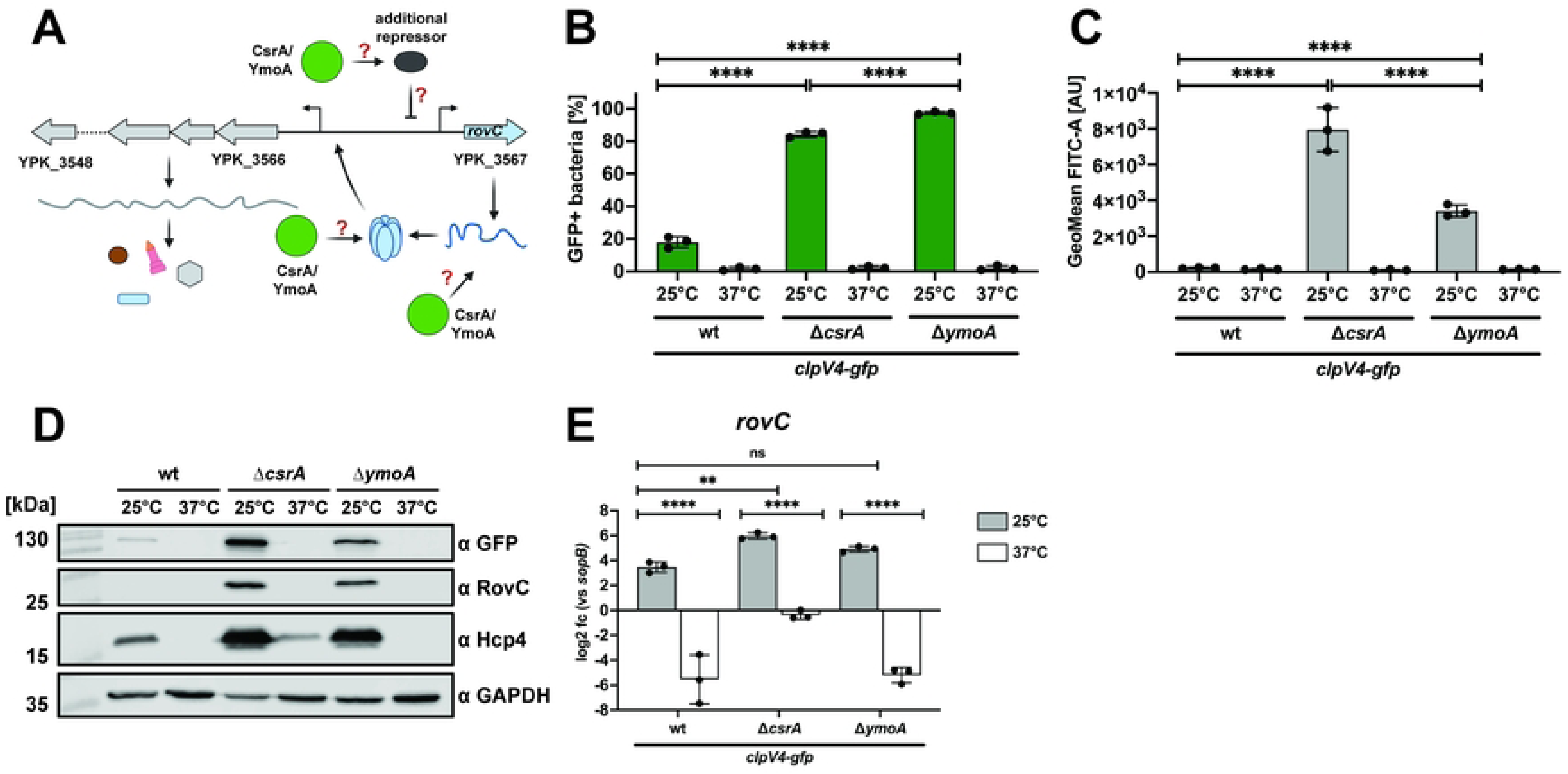
Influence of CsrA and YmoA on the temperature-controlled RovC and T6SS4 components synthesis. (A) Schematic overview of potential regulation pathways, controlling the temperature-dependent synthesis of RovC and T6SS4. Regulation can be mediated by an independent repressor or by affecting transcript or protein stability on a post-transcriptional level. Created in BioRender. Dersch, P. (2025) https://BioRender.com/9cztzkr. (B-C) Quantification of *clpV4-gfp* expressing bacteria (B) and GFP expression intensity (GeoMean FITC-A) (C) using the wt *clpV4-gfp* (YP412), Δ*csrA clpV4-gfp* (YP559), and Δ*ymoA clpV4-gfp* (YP474) strains incubated overnight at either 25°C or 37°C. 1 x 10^5^ bacteria were analyzed using flow cytometry. Data depict the mean and standard deviation of three independent experiments. Significant differences in B) and C) were determined using a Two-Way ANOVA with Tukey’s correction. (D) Western blotting of samples analyzed in (B). ClpV4-GFP, RovC, and Hcp4 protein levels were detected using antibodies against GFP, RovC, and Hcp4. RNAP was detected as a loading control. Experiments were performed in three independent replicates; one representative Western blot is shown. (E) Total RNA of an overnight culture of wt, Δ*csrA* (YP53), and Δ*ymoA* (YP50) was isolated to perform qRT-PCR. Specific primer pairs for *rovC* were used to determine the expression of *rovC*. Log2 fold changes were calculated between *rovC* and *sopB* as a non-temperature-regulated reference gene [6]. Experiments were performed in three biological replicates, and significant differences were determined using a Two-Way ANOVA with uncorrected Fisher’s LSD. ** = p ≤ 0.01, **** = p ≤ 0.0001, ns = not significant, p > 0.05.

### Downregulation of T6SS4 expression at 37°C is promoted by post-transcriptional control of *rovC* mRNA levels

A previous experiment in this study comparing *rovC* mRNA levels at different temperatures revealed that the amount of *rovC* transcripts is significantly reduced at a growth temperature of 37°C compared to 25°C (**Fig 3E**, **Fig 4E**). To gain further insight into the mechanism underlying temperature control, we tested the expression of the *rovC* gene in response to temperature using a translational reporter fusion (p*P_rovC_rovC’-‘mCherry*). As shown in **Fig 5A**, no major shifts of the *rovC-mCherry*^high^ and *rovC-mCherry*^low^ populations were detectable after 6 h of incubation at 37°C. Although similar numbers of *rovC-mCherry*^high^ expressing bacteria were detectable 6 h after the temperature upshift, we could only detect *clpV4-gfp* (T6SS4-ON) expressing bacteria in cultures grown at 25°C (**Fig 5A, B, S 5 Fig.**), indicating that the activity of the *rovC* promoter is not subjected to temperature control.

**Fig 5.**
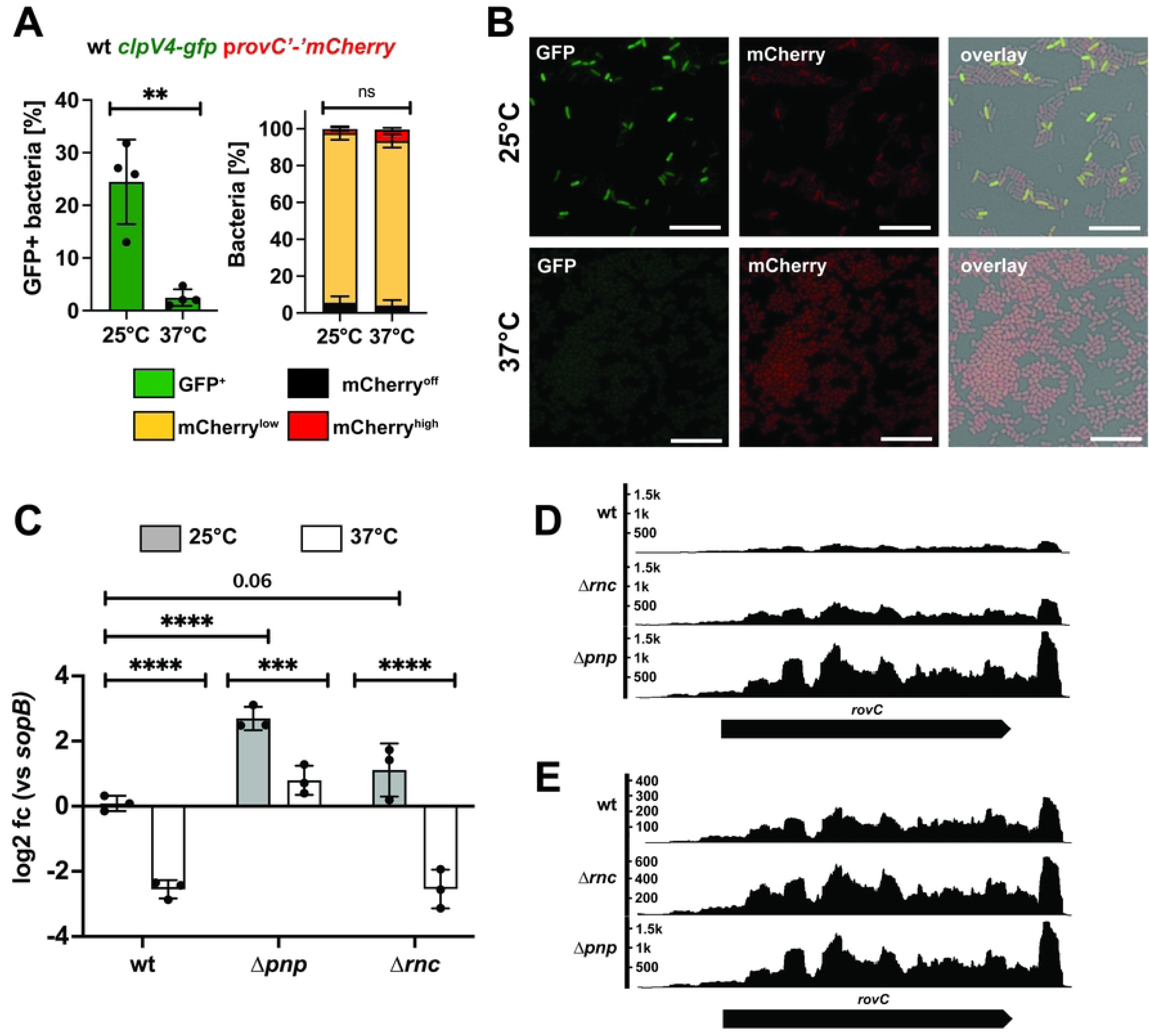
*rovC* mRNA transcript is post-transcriptionally modified at 37°C. (A) Wt *clpV4-gfp* expressing a plasmid-encoded P*_rovC_ rovC*’-‘*mCherry* fusion was incubated for 6 h at 25°C or 37°C, and *gfp-* and *mCherry*-expressing bacteria were quantified by flow cytometry. 1 x 10^5^ bacteria were analyzed for each time point using flow cytometry. Experiments were performed in four independent replicates; the mean and standard deviation are shown. Significant differences of GFP^+^ bacteria at 25°C and 37°C were determined using an unpaired t-test and of *mCherry*-expressing bacteria using a Two-Way ANOVA with Tukey’s correction. (B) Fluorescence microscopy of wt *clpV4-gfp provC’-‘mCherry* after 6 h of incubation at 25°C or 37°C. Representative images of the GFP and mCherry channels and an overlay of the brightfield and the GFP and mCherry channels were shown. Bacteria were imaged on agarose pads containing 1% agarose, and the scale bar represents 10 µm. (C) qRT-PCR was performed with total RNA extracted from wt, Δ*pnp*, and Δ*rnc* incubated for 2 h at 25°C or 37°C. Specific primer pairs were used to determine expression levels of the *rovC* gene, and log2 fold changes to *sopB* as a non-temperature-regulated reference gene [6] were calculated. Experiments were performed in three independent replicates, and significant differences were determined using a Two-Way ANOVA with Tukey’s correction. (D-E) RNA coverage of the *rovC* gene of wt, Δ*pnp*, and Δ*rnc*, incubated for 2h at 25°C with the same scale (D) and an adjusted scale (E). Data were taken from Meyer *et al*. [46]. * = p ≤ 0.05, ** = p ≤ 0.01, *** = p ≤ 0.001, **** = p ≤ 0.0001, ****=p ≤ 0.0001, ns = not significant p > 0.05.

Based on the fact that the amount of *rovC* transcript was significantly reduced at 37°C compared to 25°C, as demonstrated by qRT-PCR (**Fig 3E**, **Fig 4E**), and by an RNA-sequencing analysis [6], we hypothesized that the *rovC* mRNA is a potential target of temperature-controlled RNase-mediated degradation. To test this, we analyzed *rovC* transcript levels in mutant strains lacking different RNase genes identified in *Y. pseudotuberculosis* YPIII [46]. We found that *rovC* transcript levels in bacteria grown at 25°C were significantly increased in mutants in which the polynucleotide phosphorylase (Δ*pnp*) or the RNase III gene (Δ*rnc*) was deleted (**Fig 5C**). The increase in *rovC* transcript levels was considerably more pronounced in the Δ*pnp* mutant compared to the Δ*rnc* mutant. We also performed a comparative *rovC* transcription profile analysis of the wildtype and both RNase mutants using an RNA-sequencing data set from our previous study [46]. As shown in **Fig 5D, E**, the overall read pattern covering the *rovC* gene is comparable, and no typical changes of the sequencing read patterns due to the processing by RNases were detectable [62–64]. This suggests that the influence of the RNases on *rovC* transcript levels is not direct. Furthermore, we revealed that the *rovC* mRNA amount was still significantly reduced at 37°C in both mutants compared to 25°C (**Fig 5C)**. Hence, both RNases control the synthesis of RovC but do not seem to be mainly involved in their temperature control.

To further investigate *rovC* transcript stability, we artificially overexpressed *rovC* encoded on a medium copy plasmid from a temperature-insensitive, arabinose-inducible promoter to exclude regulatory mechanisms influencing *rovC* transcription (**Fig 6A**). We found that RovC-overexpression under the control of the P_BAD_ promoter resulted in the synthesis of RovC at 37°C (**Fig 6B**). However, the overall amount of detectable RovC was 1.38 x higher when *rovC* was overexpressed at 25°C (**Fig 6C**). Accordingly, *rovC* transcript levels were higher at 25°C after induction (**Fig 6D**), emphasizing a post-transcriptional regulation of *rovC* mRNA levels in response to temperature. We further tested how overexpression of *rovC* under these conditions influences the induction of the T6SS4 gene cluster. For this purpose, we used a *P_T6SS4_tssA4’*-‘*gfp* reporter fusion in which the predicted T6SS4 promoter [18] and the first codons of the first gene of the operon (*tssA4*, **Fig 1A**) are fused to *gfp*. As shown in **Fig 6E** and **Fig 6F**, three hours after induction of *rovC* overexpression both, the number of GFP-expressing bacteria and the overall expression of the *gfp* reporter fusion were significantly increased at 25°C and 37°C compared to empty vector controls. In line with *rovC* transcript and RovC protein levels in the bacteria (**Fig 5B, D**), the number of T6SS-ON bacteria and the overall expression of the *P_T6SS4_tssA’*-‘*gfp* reporter fusion was still higher at 25°C compared to 37°C. This further indicates that RovC is fully functional at 37°C when overexpressed from an alternative promoter, and thermally induced changes in the protein folding and/or quaternary structure of RovC triggered by the temperature upshift do not seem to be important for T6SS4 expression control.

**Fig 6.**
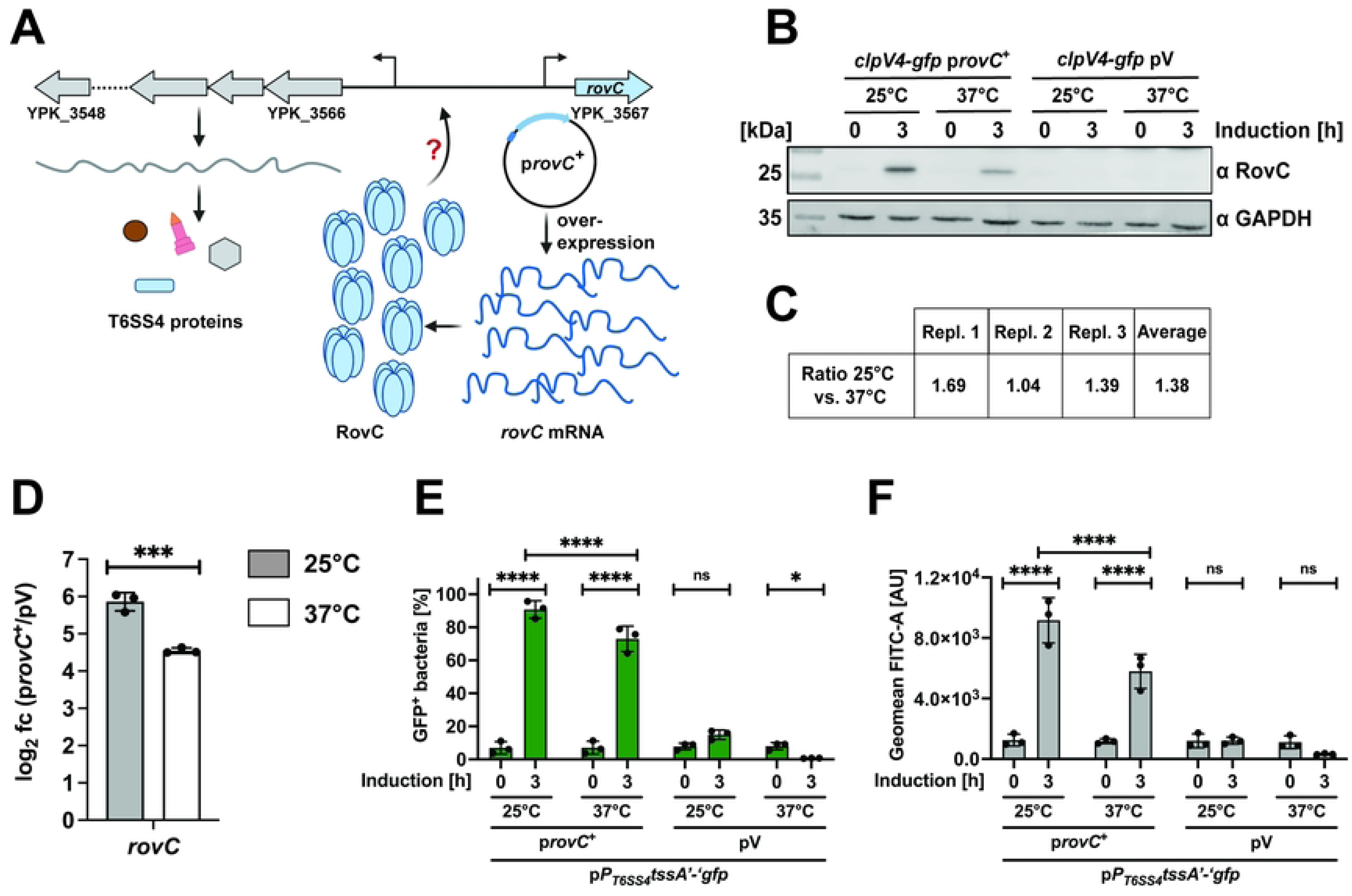
T6SS4 promoter expression can be induced by an overexpression of *rovC* at 37°C. (A) To investigate RovC functionality at 37°C, *rovC* was overexpressed from a non-temperature-sensitive P_BAD_ promoter (P_BAD_ *rovC*, p*rovC*^+^). Created in BioRender. Dersch, P. (2025) https://BioRender.com/vt98mb7. (B) Western blotting of wt *clpV4*-*gfp* harboring P_BAD_ *rovC*^+^ (p*rovC*^+^) or empty vector (pV) 0 h and 3 h after inducing overexpression of *rovC* at 25°C or 37°C. Protein levels of RovC were detected with a specific antibody against RovC, and GAPDH was used as a loading control. Experiments were performed in three independent replicates; one representative Western blot is shown. (C) Ratio of RovC between 25°C and 37°C. The amount of detected RovC of all three Western blot replicates was normalized to the loading control, and the ratio between 25°C and 37°C was calculated. (D) Total RNA of samples in (B) was extracted to perform qRT-PCR to determine expression levels of the *rovC* gene. Expression levels were normalized to *sopB* as a non-temperature-regulated reference gene [6] and log2 fold changes 3 h after inducing overexpression of *rovC* between p*rovC*^+^ and empty vector control (pV) were calculated. Experiments were performed in three independent replicates. Significant differences were determined using an unpaired t-test. (E-F) Promoter region of T6SS4 and first codons of *tssA4* were translationally fused to *gfp* on a plasmid to identify bacteria with an active T6SS4 promoter by flow cytometry. Strains additionally harbored p*rovC*^+^ or empty vector (pV), and overexpression of *rovC* was induced for 3 h at either 25°C or 37°C. The amount of GFP^+^ bacteria (E) and the expression intensity (F) of 1 × 10^5^ bacteria were analyzed, and the data depict the mean and standard deviation of three independent experiments. Significant differences were determined using Two-Way ANOVA with Tukey’s correction. * = p≤0.05, *** = p ≤ 0.001, **** = p ≤ 0.0001, ns = not significant p > 0.05.

### Processing of the T6SS4 polycistronic mRNA leads to differential expression of T6SS4 genes in response to temperature

In our attempt to analyze how RovC influences T6SS4 transcript levels at different temperatures, we also tested the expression of the translational *clpV4-gfp* fusion after *rovC* overexpression (**Fig 7A**). Even though the P_T6SS4_ promoter was highly induced at 37°C in approximately 70% of the bacteria (**Fig 6E**), we could not detect an equivalent induction with the *clpV4-gfp* fusion. Compared to 25°C with 90% GFP-positive bacteria (T6SS4-ON), less than 5% of the population could be identified as T6SS4-ON at 37°C (**Fig 7B**). This suggests that not only the level of *rovC* mRNA but also that of T6SS4 transcripts is influenced by temperature. A qRT-PCR analysis comparing the transcript levels of five T6SS4 genes (*tssA4, vipA4, hcp4, clpV4,* and *tssK4*) after RovC induction supported this assumption (**Fig 7C**). The abundance of most transcripts was significantly lower at 37°C compared to 25°C (**Fig 7C**). One exception is *hcp4,* for which similarly high levels of the *hcp4* transcript and the Hcp4 protein were detected at 25°C and 37°C temperatures (**Fig 7C**, **D**).

**Fig 7.**
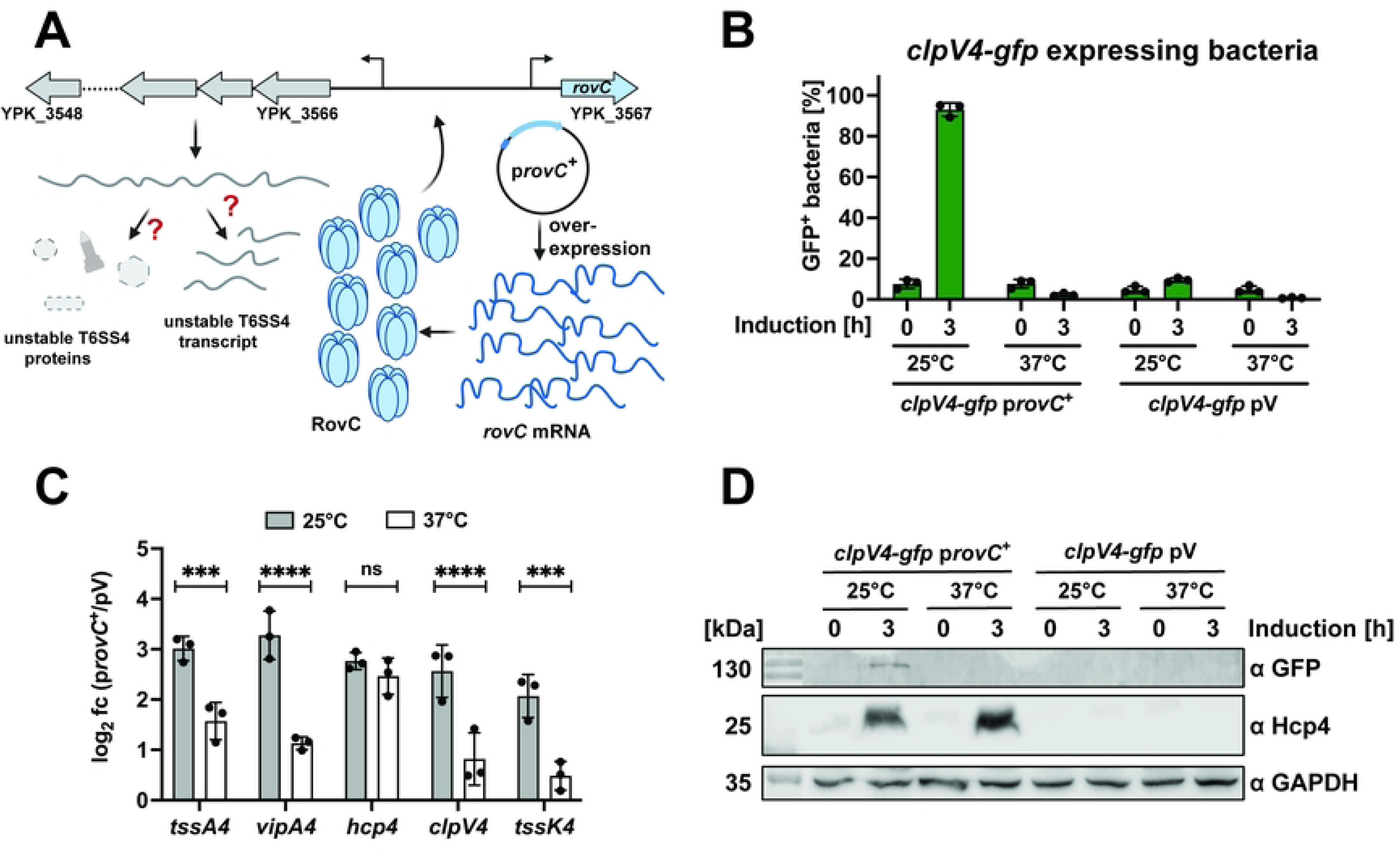
The overall amount of individual T6SS4 gene mRNAs differs significantly at 37°C and 25°C. (A) Scheme of potential downregulation of T6SS4 expression at 37°C caused by an unstable T6SS4 transcript or T6SS4 proteins. Created in BioRender. Dersch, P. (2025) https://BioRender.com/947o0tl. (B) Quantification of *clpV4-gfp* expressing bacteria after inducing overexpression of *rovC* for 3 h at either 25°C or 37°C. Data depict the mean and standard deviation of three independent experiments. (C) Total RNA of samples analyzed in B) was extracted for qRT-PCR. Specific primer pairs for five T6SS4 genes were used to determine gene expression within the T6SS4 operon. All expression levels were normalized to *sopB* as a reference gene, and log2 fold changes 3 h after overexpressing *rovC* between p*rovC*^+^ and empty vector control (pV) were calculated. Experiments were performed in three independent replicates, and significant differences were determined using a Two-Way ANOVA with Šidák correction. (D) Western blotting of the same samples analyzed in (B). Protein levels of ClpV4-GFP and Hcp4 were detected using specific antibodies against GFP and Hcp4. GAPDH was used as a loading control. One representative Western blot out of three independent experiments is shown. * =p ≤ 0.05, *** =p ≤ 0.001, **** =p ≤ 0.0001, ns = not significant p>0.05.

It is unlikely that this expression pattern is caused by an additional promoter located upstream within the T6SS4 operon, as no promoter or an additional transcriptional start site upstream of *hcp4* is predicted [6,18]. Moreover, a translational *hcp4’-‘gfp* fusion construct harboring the 5’UTR upstream of *hcp4* without the T6SS4 promoter was not expressed (**S 6 Fig.**). Taken together, this suggests that in addition to the *rovC* mRNA, the transcripts of the individual T6SS4 genes are post-transcriptionally controlled, but to a different extent.

To prove this, we compared the transcript levels of eight genes covering different regions of the T6SS4 operon under native (non-RovC-inducing) conditions using qRT-PCR (**Fig 8A, B**). We found that the overall abundance of transcripts covering the different genes of the operon varied significantly. The transcript levels of the first four encoded genes in the operon (*tssA4, vipA4, vipB4, hcp4*) were significantly higher compared to the genes further downstream (*tssE4, tssF4*, *clpV4, tssK4*) (**Fig 8B**). A comparison of the qRT-PCR data with the transcription profile of the individual T6SS4 genes using our RNA-sequencing data supported this observation and confirmed a particularly high abundance of the *hcp4* transcript (**S 7 Fig.**).

**Fig 8.**
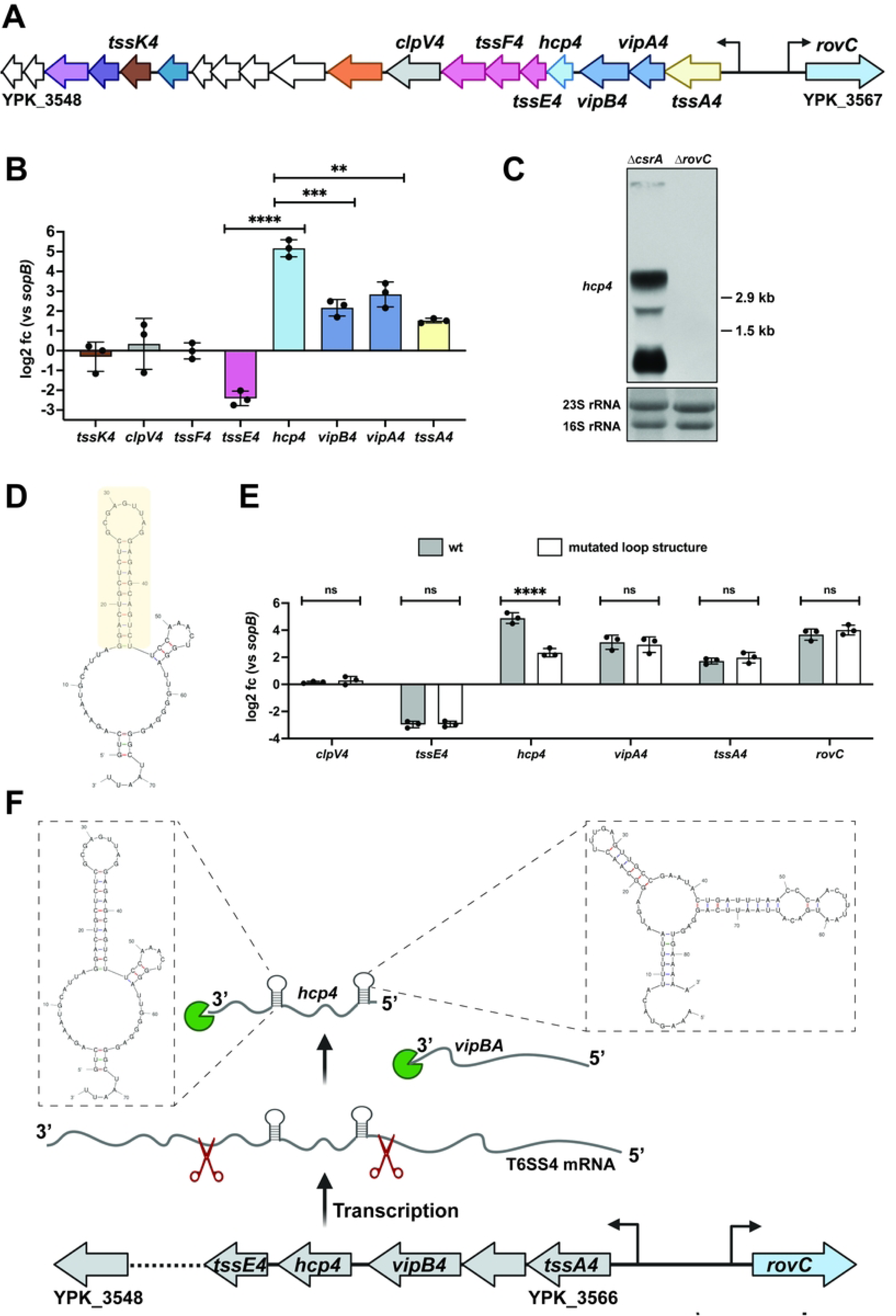
Different abundance of individual T6SS4 gene transcripts. (A) Schematic overview of the T6SS4 operon. (B) Total RNA of *Y. pseudotuberculosis* wt incubated for 6 h at 25°C was extracted to perform qRT-PCR. Specific primer pairs for eight T6SS4 genes were used to determine expression levels within the operon, log2 fold changes to *sopB* as a non-temperature-regulated reference gene [6] were calculated. Experiments were performed in three independent replicates, and significant differences were determined using an unpaired t-test. (C) Northern blot of total RNA of YP53 (Δ*csrA*) and YP154 (Δ*rovC*). A specific probe for *hcp4* was used; 16S and 23S rRNA were used as loading controls. (D) Secondary structure of the wildtype intergenomic region between *hcp4* and *tssE4* was predicted using the mFold web server [65]. The region which was mutated is marked in yellow. (E) Total RNA of *Y. pseudotuberculosis* wt and a strain with the mutated loop structure grown for 6 h at 25°C was extracted to perform qRT-PCR. Specific primer pairs for six T6SS4 genes were used to determine expression levels within the operon, and log2 fold changes to *sopB* as a non-temperature-regulated reference gene [6] were calculated. Experiments were performed in three independent replicates, and significant differences were determined using a Two-Way ANOVA with Šidák correction. (F) Scheme of potential post-transcriptional T6SS4 regulation. The T6SS4 transcript is processed due to endoribonucleolytic cleavage (red scissors). The 5’ and 3’ protection loop up- and downstream of *hcp4* protects the *hcp4* transcript from further endonuclease activity and from being degraded by exoribonucleases (green pie). The loop structures of the 5’-UTR and 3’-UTR of *hcp4* were predicted using the mFold web server [65]. Created in BioRender. Dersch, P. (2025) https://BioRender.com/efeis70. ** = p≤0.01, *** = p≤0.001, **** = p ≤ 0.0001, ns = not significant p > 0.05.

Subsequent Northern blotting with an *hcp4* probe revealed multiple bands of different sizes, further indicating a processed T6SS4 transcript (**Fig 8C**). As T6SS4 expression in the *Y. pseudotuberculosis* wt is generally very low, the *csrA* mutant was used for better visualization of the *hcp4* transcript on the Northern blot, as deletion of this global regulator leads to an overall upregulation of T6SS4 gene cluster expression (**Fig 4**, **S 4 Fig.**). A *rovC* mutant strain was used as a negative control, as this abolishes T6SS4 expression [30].

The results strongly suggest that the differences in T6SS4 transcript levels result from post-transcriptional control mechanisms such as differential mRNA degradation, stabilization, and/or transcriptional termination. Secondary structure prediction of the intergenomic region between *hcp4* and *tssE4* revealed a GC-rich stem loop directly downstream of the *hcp4* gene (**Fig 8D**). The hairpin stem was chromosomally mutated to test whether this loop functions as a potential transcription terminator, which would explain the observed transcriptional stop after *hcp4* (**Fig 8D**, marked in orange). qRT-PCR revealed no differences in the transcript levels of the tested T6SS4 genes (**Fig 8E**). Only the transcript level for *hcp4* was significantly reduced in the strain with the mutated stem loop compared to wt. This indicates that the loop may not function as a transcriptional terminator to pause transcription downstream of *hcp4*, but rather acts as a 3’-protective loop to resist exoribonucleolytic degradation of *hcp4* mRNA after endolytic cleavage at this site (**Fig 8F**). The inspection of the T6SS4 cluster further revealed another potential stem-loop structure that could be formed in the intergenic region between *vipB4* and *hcp4* (**Fig 8F**). The intergenomic region additionally consists of several AUUA-motifs, which could be recognized and then cleaved by endoribonucleases. As the *hcp4* probe interacted with a small RNA fragment of the expected size of the *hcp4* gene of around 500 bp (**Fig 8C**), it is likely that this stem-loop structure also contributes to processing and/or stabilization of the *hcp4* transcript.

## Discussion

*Y. pseudotuberculosis* is a widespread environmental bacterium that can also infect mammals [1,2]. This dual lifestyle requires a precise and rapid change in gene expression in response to changing conditions, such as entry into the human body. In this context, temperature plays an important role in the regulation of gene expression in *Y. pseudotuberculosis*, since many virulence-associated and metabolic genes are strictly controlled by temperature [5,6]. In this study, we could show that the T6SS4 in *Y. pseudotuberculosis* is strongly regulated by temperature but in the opposite manner to the T3SS. At moderate temperatures (25°C), the T6SS4 gene cluster is heterogeneously expressed, with only 10-15% of the population producing the transcriptional regulator RovC at levels which are sufficient to induce T6SS4 expression. At 37°C, *rovC* is rapidly downregulated on post-transcriptional level, leading to a rapid and complete shut-off of T6SS4 gene expression. Moreover, T6SS4 is part of a complex regulatory network that is embedded in global regulation. We found that global regulators such as the carbon starvation system regulator CsrA, the global protease Lon, the *Yersinia* modulator protein YmoA, and the two RNases PNPase and RNase III repress RovC and T6SS4 synthesis at 25°C. This indicated a very sophisticated and tight control of T6SS4 gene expression in response to temperature and other environmental signals that differs fundamentally from the control of T6SS expression in other bacteria.

In this study, a deeper analysis of T6SS4 expression in this study showed that differential expression levels of the *Yersinia*-specific transcriptional activator RovC drive T6SS4 heterogeneity (T6SS4-ON versus T6SS4-OFF) (**Fig 1**, **Fig 2**). Phenotypic heterogeneity has been intensively studied in recent years and can be triggered, for example, by microenvironments with different demands, leading to different gene expression within a subpopulation [66,67]. Phenotypic heterogeneity can also be a strategy to survive sudden environmental fluctuations. In this so-called bet-hedging strategy, only some bacteria express certain genes that are not necessary under normal conditions. This additional expression of non-required genes can cost energy and resources. Nevertheless, since no adaptation phase is necessary in the event of sudden changes, the survival of at least a subpopulation is ensured [68–72]. Such bistability of gene expression has been demonstrated for *rovA* in *Y. pseudotuberculosis* controlling invasin expression in response to temperature [73–75]. RovA undergoes a conformational change upon an upshift from 25°C to 37°C that increases its accessibility for proteases, and hence, its rapid degradation abolishes its positive autoregulation (positive feedback loop) [73–75]. The exact mechanism of how heterogeneity of *rovC* and thus, T6SS4 expression is achieved is still unclear. However, no autoregulation or -activation of RovC was detectable [30], indicating a substantially different mechanism from RovA. Moreover, the functional role of T6SS4 remains unknown. However, the presence of four different T6SS clusters in *Y. pseudotuberculosis* likely requires tight regulation to ensure that T6SS4 expression is restricted to the environments where it is most advantageous.

In contrast to *Y. pseudotuberculosis*, T6SS is not heterogeneously but equally expressed in most studied bacteria, including *Vibrio* and *Pseudomonas,* where the secretion system genes have been extensively studied [19,22,23,26,48–50]. Only one recent study in enteroaggregative *E. coli* (EAEC) has reported that its T6SS is ON in 60% and OFF in 40% of the bacteria [76]. Yet, the implicated regulatory factors in EAEC are very distinct from *Y. pseudotuberculosis.* They found that heterogeneous T6SS expression in EAEC is regulated by the interaction of Fur (ferric uptake regulator) with the T6SS promoter region and genetically controlled by GATC methylation sites [76]. In contrast, heterogeneous T6SS4 expression in *Y. pseudotuberculosis* is already determined by the heterogeneous expression of the transcriptional regulator RovC (**Fig 2**). In addition, neither Fur binding sites nor GATC methylation sites have been found for RovC, suggesting a different regulation of heterogeneous T6SS4 expression *in Y. pseudotuberculosis*.

Our study further showed that RovC synthesis is the major control hub for T6SS4 expression. All global regulators of T6SS4 and temperature changes seem to influence *rovC* transcript levels strongly but do not affect its promoter activity or protein stability. Several regulatory scenarios are possible. For instance, differential folding of the mRNA in response to temperature could expose or hide RNase cleavage sites [77,78] and/or influence translation efficiency [77,79–85]. Alternatively, the synthesis or activity of RNases targeting the *rovC* mRNA could be increased at higher temperatures, leading to degradation of the *rovC* transcript.

Like other functionally linked genes, all core components of T6SS4 are encoded in close proximity under the control of only one promoter and form a polycistronic mRNA [18]. As a result, only one mRNA is transcribed, which, in theory, should lead to stoichiometrically equal amounts of transcripts. However, not all components are needed to the same extent, and regulatory measures are implemented to adjust T6SS4 component synthesis to their functional need. In the case of the T6SS, for example, proteins that assemble the membrane or the base plate complex are required in lower quantities than those building the tubular structure, which consists of hundreds of copies of stacked Hcp proteins [21,86]. While different translation efficiencies have been described to resolve this problem in some cases [87,88], we show that T6SS4 gene expression differences have already occurred at the transcript level. Very low transcript levels were detected for the genes downstream of *hcp4.* In contrast, the mRNA abundance of the first four genes of the operon was significantly higher, particularly *hcp4*, encoding the tube protein. No additional promoter activity was detected upstream of *hcp4* (**S 6 Fig.**), and no other additional transcriptional start site within the operon has been identified [6,18]. Thus, differential T6SS4 mRNA levels seem to result from endoribonucleolytic cleavage of the polycistronic T6SS4 cluster mRNA at two sites with stem-loop structures flanking the *hcp4* gene. A very rapid exoribonucleolytic degradation can explain the low abundance of the T6SS gene transcripts downstream of *hcp4*. In contrast, both stem loops flanking the resulting short *hcp4* transcript will likely protect the transcript from further cleavage, leading to very high *hcp4* transcript and thus Hcp4 protein levels (**Fig 8C**, **Fig 5D**). Differential processing and degradation of polycistronic mRNA are described for several bacterial operons [77,77,89–93]. Processing of polycistronic mRNAs, resulting in stabilization of upstream genes (*e.g.,* by the presence of stem loop structures, as identified downstream of *hcp4*) was described for the maltose and *iscRSUA* operon in *E. coli* or the *puf* operon in *Rhodobacter capsulatus* [77,89,90,94]. Endonucleolytic cleavage of mRNAs can also lead to a lower abundance of upstream transcripts (as observed for *tssA4*, *vipA4, vipB4*, compared to *hcp4*) due to degradation by 3’-5’-exoribonucleases. Such a mechanism has been demonstrated for the *pap* and glycogen operon in *E. coli* [77,91,92,95]. Alternatively, as reported in other systems [89,96,97], strong translation of the *hcp4* gene could prevent this gene from being degraded by 3’-5’ exoribonucleases.

The observed post-transcriptional regulation of the T6SS4 polycistron, leading to a different abundance of T6SS4 transcripts, fits with studies reporting differential T6SS protein amounts [86]. Some proteins, such as those coding for the baseplate, require only a small number of copies, whereas the tube and sheath structure components require many copies [21,29,52,98–100]. After a firing event, the sheath structure components VipA and VipB are disassembled by the ATPase ClpV and remain in the cell [29,49,101,102]. In contrast, Hcp is secreted with VgrG and the effector proteins into the target organism and can be found in the supernatant after a firing event [21,26,103,104]. This requires immediate *de novo* synthesis of the secreted proteins in case of a new firing event. To solve this, other organisms such as *Vibrio spp*. encode, in addition to the main T6SS island, auxiliary clusters with *hcp* or effector proteins under the control of a separate promoter [20,50,105,106]. Since a single promoter controls all T6SS4 genes in *Y. pseudotuberculosis*, the observed differential stabilization of the respective mRNAs is an effective way for *Y. pseudotuberculosis* to meet the different protein requirements.

To date, the precise role of T6SS4 is still unclear. The highest expression of *rovC* and T6SS4 at moderate temperatures and their strong repression at 37°C observed in this and other studies (**Fig 1C**, **Fig 5A**, [18,31]) indicate a function outside mammalian hosts. This is supported by multiple RNA sequencing datasets, including an *in vivo* transcription profile of *Y. pseudotuberculosis* within the Peyer’s patches during a mouse infection [6,44,46,47], revealing a complete repression of T6SS4 gene expression at body temperature. Most interestingly, we found T6SS4-type clusters in the environmental-associated strains *Winslowiella toletana, Serratia fonticlola, Enterobacillus tribolii, Trabulsiella guamensis, T. odontotermitis,* which belong to the family of *Enterobacteriaceae* (**Fig 9**). The identified T6SS clusters are homologous in synteny and protein identity to the T6SS4 of *Y. pseudotuberculosis* and T6SS-A of *Y. pestis.* Among them, *W. toletana* shows the highest protein identity to the T6SS4 of *Y. pseudotuberculosis,* including the gene of the so far unique transcriptional activator RovC (**Fig 9**, > 70% amino acid identity). Initially described as *Erwinia toletana*, *W. toletana* exhibits an optimal growth temperature between 28°C and 30°C [107,108]. It was first isolated from olive knots in association with the plant pathogen *P. savastanoi*, and its presence has been shown to enhance olive tree infection [107,109]. *T. guamensis* was found in soil, vacuum cleaner dust, or human stool samples, *T. odontotermitis* and *E. tribolii* were isolated from the gut of termites and red fluor beetle, respectively [110–113]. *S. fonticola* is a widely distributed environmental bacterium, commonly found in soil or aquatic habitats [114,115]. Despite a few clinical cases of *S. fonticola*, none of the organisms are considered primary human pathogens. The high amino acid identity, particularly for *W. toletana* strongly indicates a similar function of the T6SSs, possibly linked to a plant- or insect-associated ecological niche [116].

**Fig 9.**
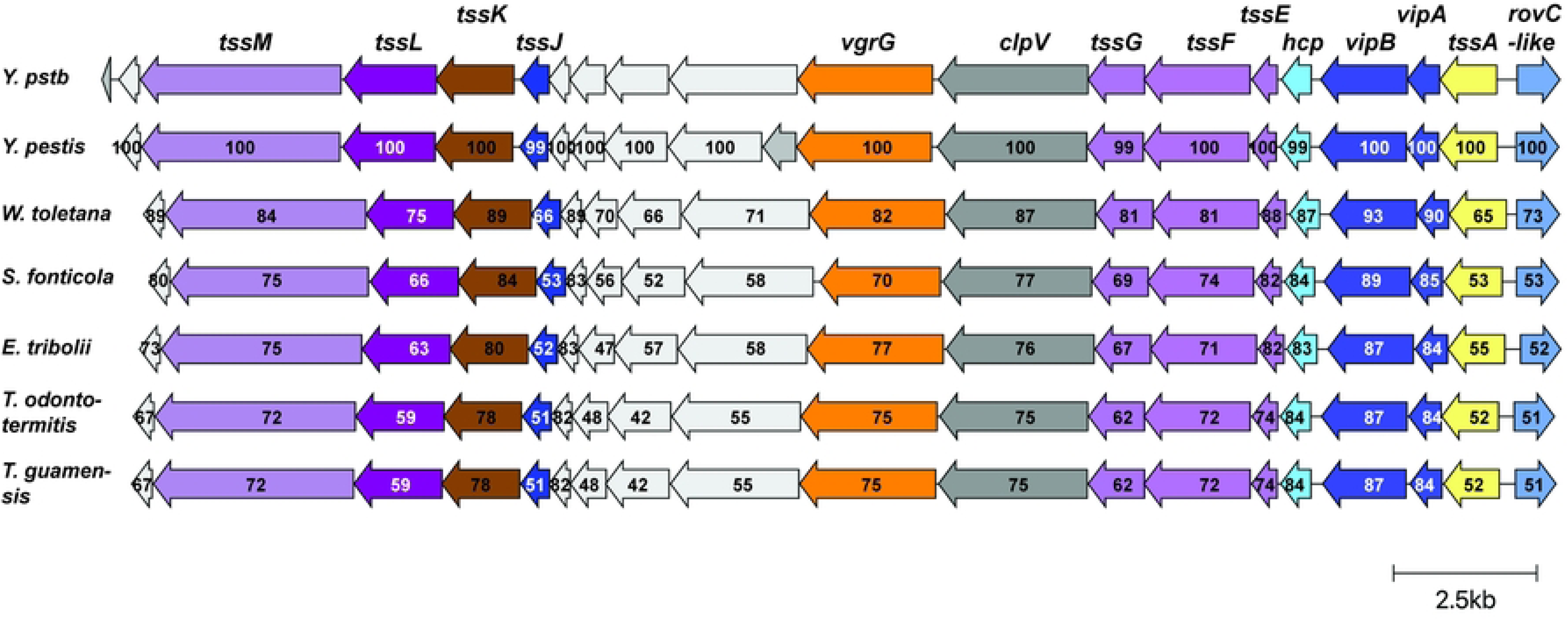
Identification of homologous T6SS4 cluster in different *Enterobacteriaceae*. The amino acid identity of T6SS4 components between *Y. pseudotuberculosis* and different strains is given as a percentage based on protein sequence similarity. Visualization of the cluster similarities was done using clinker webserver [117]. The scale bar represents 2.5 kb.

In contrast to these assumptions, one group reported that some potential T6SS4 effectors are essential for virulence in mice [40,42,43]. One of these potential effectors (YPK_3548) is encoded at the 3’-end of the T6SS4 cluster. This gene is however not present in the T6SS4-type cluster in *Y. pestis* and *W. toletana* (**Fig 9**), and its expression was not affected by the loss of the *csrA* gene, in contrast to all other T6SS4 cluster genes [30]. As all *in vitro* experiments to characterize the effectors were also carried out at 26°C [40,43], a more detailed analysis of the *in vivo* function of the T6SS4 system and its putative effectors is required to elucidate whether the T6SS4 system is important for a plant- and/or animal-associated lifestyle.

## Material and Methods

### Media and growth conditions

*E. coli* was grown overnight in 5 ml Luria-Bertani (LB) medium (5 g/l yeast extract, 10 g/l tryptone, 5 g/l NaCl) at 37°C. Super Optimal Broth with Catabolic Repression (SOC) medium (5 g/l yeast extract, 20 g/l tryptone, 10 mM NaCl, 2.5 ml KCl, 10 mM MgSO_4_, 10 mM MgCl_2_, 20 mM glucose) was used to allow *E. coli* cells to recover after transformation. The medium is based on Super Optimal Broth (SOB) medium [118], supplemented with glucose. *Y. pseudotuberculosis* was grown in LB (BD Bioscience, USA), supplemented with 1 mM CaCl_2_ or in brain-heart infusion (BHI) medium (37 g/l BHI, BD Bioscience, USA). Unless stated otherwise, *Y. pseudotuberculosis* cultures were incubated overnight at 25°C in LB, main cultures at either 25°C or 37°C. If necessary, selective antibiotics were added in the following final concentration: 100 µg/ml ampicillin, 50 µg/ml kanamycin, and/or 50 µg/ml chloramphenicol. To induce overexpression of *rovC* from pBAD30 plasmid, 0.1% arabinose was added to the culture in early exponential phase (OD_600_ 0.7).

### Plasmid and strain construction

In this study, molecular cloning of DNA into vectors was performed following the Gibson Assembly Protocol (E5510) (New England Biolabs), based on the method developed by Gibson *et al*. [119]. Plasmid DNA was isolated using the Nucleospin Plasmid kit (Macherey Nagel), and genomic DNA of *Y. pseudotuberculosis* was isolated using the ISOLATE II Genomic DNA Kit (Bioline). The oligonucleotides used for cloning, sequencing and qRT-PCR were purchased from Eurofins Genomic. PCRs for fragment amplification were performed using Q5 High-Fidelity 2X Master Mix (New England Biolabs) following the manufacturer’s instructions. PCR products were purified using Nucleospin Gel and PCR clean-up kit (Macherey Nagel). To verify correct clones, colony PCRs were performed using 2x DreamTaq Green PCR Master Mix (Thermo Scientific) according to the manufacturer’s instructions. Sanger sequencing (Microsynth Seqlab GmbH) confirmed successful cloning of plasmids. Plasmids and primers used in this study are listed in **Table S 1** and **Table S 2**.

For cloning of pANK4 and pANK15, vector pFU98 was linearized by digestion with *Nhe*I and *Not*I (New England Biolabs, Ipswich, USA). For pANK4, the 5’-UTR of *rovC* (-579 to +13) was amplified with primers IX691/692 using Q5 High-Fidelity 2X Master Mix (New England Biolabs). Primers IX687/IX688 were used to amplify *mCherry*. A splicing by overlap extension PCR (SOE PCR) [120] was performed to generate a combined DNA fragment of 5’-UTR of *rovC* and *mCherry*. The combined DNA fragment was cloned into linearized pFU98 following the Gibson Assembly Protocol (E5510) (New England Biolabs). For generating pANK15, primers X108/X109 were used to amplify the 5’-UTR of YPK_3566, including the predicted promoter region of T6SS4 (-581 to +15) [18]. Primers X110/111 were used to amplify *gfpmut3.1*. SOE PCR was performed to generate the combined DNA fragment with primers X108/X111, which was subsequently cloned into linearized pFU98. For cloning of pANK25, vector pFU31 was linearized with *Sal*I and *Nhe*I (New England Biolabs, Ipswich, USA). The 5’UTR of *hcp4* (-212 to +30) was amplified with primers X382/X383. The amplified DNA fragment was subsequently cloned into linearized pFU31. For cloning of pANK45, the suicide plasmid pAKH3 was linearized by digestion with *Xma*I and *Sph*I (New England Biolabs, Ipswich, USA). The upstream and downstream region of the intergenomic region between *hcp4* (YPK_3563) and *tssE4* (YPK_3564) were amplified with primers X558/X559 and X560/X561, respectively. A combined DNA fragment of the upstream and downstream region was generated with primers X558/X561 performing a SOE PCR.

All generated vectors were transformed into electrocompetent *E. coli* SM10 λpir. 500 ml of LB were inoculated 1:100 with an *E. coli* overnight culture and grown to an OD_600_ of 0.6 at 37°C. The culture was pelleted for 10 min at 8,000x g and 4°C, followed by two washing steps with 50 ml and 25 ml ice-cold water. After an additional washing step in 1 ml ice-cold water, the cells were centrifuged again for 10 min at 8,000 x g and 4°C. The pellet was resuspended in a final volume of 1.8 ml ice-cold 10% glycerol. 50 µl aliquots were immediately frozen in liquid nitrogen and stored at -80°C. For transformation, 2 µl of plasmid DNA was added to 50 µl competent cells and exposed to 2.2 kV (200 Ω, 25 μF) for 5 ms in a pre-cooled electroporation cuvette. Transformed *E. coli* cells were mixed with 1 mL SOC and incubated for 1 h at 37°C with permanent aeration to recover. Afterwards, the cells were pelleted for 2 min at 8,000 x g, resuspended in 100 µl medium, plated out, and incubated overnight at 37°C on agar plates with the desired antibiotic. Correct cloning of generated plasmids was confirmed by Sanger sequencing.

Electrocompetent *Y. pseudotuberculosis* strains were used to transform pVK25, pBAD30, pANK4, pANK15, pANK25, pFU31, pKD4, and pCP20. The 15 ml BHI medium was inoculated 1:50 with *Y. pseudotuberculosis* overnight cultures. After 3 h of incubation at 25°C, the culture was pelleted for 5 min at 8,000 x g and 4°C and washed twice with 5 ml ice-cold sterile water. After the second wash, the pellet was resuspended in 200 µl ice-cold water and immediately used for transformation. For each transformation, 2 µl plasmid DNA and a 50 µl aliquot of competent cells were added to a pre-cooled electroporation cuvette. Transformation was performed by exposing the cells for 5 ms to 2.2 kV (200 Ω, 25 μF). Subsequently, the transformed cells were allowed to recover in 1 ml BHI medium for 2 h at 25°C under constant aeration. 100 µl of each transformation was plated on agar plates with the corresponding antibiotics and incubated at 25°C for two days.

### Construction of *Y. pseudotuberculosis* mutants

All strains used in this study are listed in **Table S 1**. Chromosomal *Y. pseudotuberculosis* mutants were generated by homologous recombination using suicide plasmids derived from pAKH3 [121], except for YP78. For chromosomal fusion of *clpV4* to *gfp,* pASS90 was conjugated into the respective *Y. pseudotuberculosis* strains. For chromosomal deletion of the intergenomic region between *hcp4* and *tssE4*, pANK45 was conjugated into *Y. pseudotuberculosis* wildtype [122]. Conjugation of *Y. pseudotuberculosis* was followed by sucrose selection on agar plates containing 6% sucrose. After conjugation, correct clones were confirmed by PCR and Sanger sequencing (Microsynth Seqlab GmbH).

To chromosomally delete *lon* (YPK_3232), a Red-mediated recombination method was used as described [123]. In brief, primers I407/I408 were used to amplify a kanamycin resistance cassette with homologous regions of YPK_3232 using pKD4 as template. The amplified DNA fragment was transformed into YPIII harboring pKD4. Loss of pKD4 and successful gene disruption were confirmed with colony PCR. The kanamycin resistance cassette was eliminated by transformation of the mutated strain with pCP20. Afterwards, the plasmid was cured by incubating for 3h at 42°C due to the temperature-sensitive replicon [123,124].

### Fluorescence-based flow cytometry

Flow cytometry was performed as a high-throughput method to analyze gene expression and promoter activity using the CytoFLEX S (Beckmann Coulter, USA). To avoid the detection of spillover signal, a compensation matrix was applied for all used fluorochromes according to the manufacturer’s manual. To separate bacteria from the debris, the samples were stained with 5 µg/ml 4′,6-diamidino-2-phenylindole (DAPI) and detected by the PB450 channel. The ECD channel was used to differentiate live and dead cells, which were stained with 1 µg/ml propidium iodide (PI). FITC was used to detect GFP-expressing bacteria. In experiments, where *rovC* promoter activity was analyzed by gating for mCherry^+^ bacteria, the samples were not stained with PI, and mCherry^+^ bacteria were analyzed in the ECD channel. The threshold was set to 2500, defined by sideward scatter height (SSC-H). In this study, the gain for the SSC was set to 400, for the forward scatter (FCS) to 100, for PB450 to 300, for ECD to 1250, and for fluorescein isothiocyanate (FITC) to 700. For the analysis of GFP- or mCherry-expressing bacteria, the samples were incubated according to the respective experiment. Depending on the OD_600_, 1-10 µl were taken directly from the culture at indicated timepoints and added to 500 µl PBS with 5 µg/ml DAPI and 1 µg/ml PI. 1 x 10^5^ bacteria were analyzed for every sample using the gating strategy displayed in **S 2.**

### Fluorescence microscopy

Fluorescence microscopy was performed using the Keyence epi-fluorescence microscope BZ-X (Osaka, Japan) at 60x magnification (Plan Apochromat 60x Oil). The samples were pelleted for 5 min at 8,000x g at room temperature and resuspended to an OD_600_ of 10. From this dense suspension, 1 µl was transferred to an agarose pad containing 1% agarose and 0.5 x PBS to avoid the swimming of motile *Y. pseudotuberculosis*. For an even surface, the agarose was poured on a microscopy slide, fitted with a gene frame (Thermo Fisher Scientific, Waltham, USA).

### RNA isolation

Bacterial cultures were grown according to the respective experiment. At indicated timepoints, 2 ml of the culture were pelleted for 2 min at 10,000x g at room temperature. The pellet was directly frozen in liquid nitrogen and stored at -80 °C until isolation of total RNA. Total RNA was extracted using the Monarch Total RNA Miniprep Kit (New England Biolabs) according to the manufacturer’s protocol: “Total RNA Purification from Tough-to-Lyse Samples (bacteria, yeast, plant, etc.)”. The first part of the sample digestion and homogenization protocol was modified as follows: After thawing, the bacterial pellet was resuspended in 200 µl TE-buffer (100 mM Tris HCl pH 7.5, 1 mM EDTA pH 8) containing 10 mg/ml lysozyme (L6876, Sigma-Aldrich) and incubated for 10 min at room temperature. Afterwards, 2x volumes RNA Lysis Buffer were added and the sample was vortexed vigorously for 10 sec. After centrifugation for 2 min at 16,000x g, the supernatant was transferred to a gDNA removal column. The extraction was proceeded with the second part of the RNA Binding and Elution protocol. Total RNA was diluted with 50 µl of nuclease-free water, and the purity and concentration were determined using the Nanodrop spectrophotometer (Thermo Fisher Scientific, Waltham, USA). Because of the high abundance of genomic DNA, an additional DNA digestion step was included after isolation of total RNA. For this, 7.5 µg RNA was filled up to 44 µl of nuclease-free water, 5 µl 10X TURBO DNase buffer, and 1 µl TURBO DNase enzyme (Invitrogen, Thermo Fisher Scientific, Waltham, USA). Samples were incubated for 30 min at 37°C, followed by adding 1 µl of TURBO DNase enzyme and incubating for 30 min at 37°C. The reaction was stopped by adding 5 µl DNase Inactivation Reagent (Invitrogen, Thermo Fisher Scientific, Waltham, USA). After inactivation, the sample was centrifuged for 3 min at 16,000 x g at room temperature, and 35 µl were transferred to a fresh reaction tube. Purity and concentration of the RNA were again analyzed using the Nanodrop spectrophotometer, and the RNA was diluted to 15 ng/µl and stored at -80°C for further experiments.

### Quantitative real-time PCR (qRT-PCR)

To determine the amount of transcript of different T6SS4 genes, qRT-PCR was performed using the Luna Universal One-Step RT-qPCR Kit (New England Biolabs) with the LightCycler 96 System (Roche). Each reaction was performed in technical triplicates in a 10 µl reaction, containing 5 µl Luna Universal One-Step Reaction Mix (2X), 0.5 µl Luna WarmStart RT Enzyme Mix (20X), 0.5 µl of each primer (10 µM) and 1 µl of the desired RNA template (15 ng/µl). The reaction was filled up with nuclease-free water to 10 µl. Relative changes in gene expression were calculated using *sopB* as a reference gene [6,125]. Primers used for qRT-PCR are listed in **Table S 2**.

### Northern blot

In order to detect the *hcp4* transcript, a DIG-labelled *hcp4*-specific probe was generated (Primers are listed **Table S 2**). Total RNA was isolated, and 15 µg/20 µl of RNA was used for the Northern blot. The samples were mixed with 4 µl of 5x loading dye (31% formamide, 2.7% formaldehyde, 0.1 mg/ml ethidium bromide, 4 mM EDTA pH 8, 20 % glycerol, 0.03% bromphenole blue, 10% MOPS buffer (20x)). The samples were boiled 2x for 10 min at 70°C with 5 min in between at 10°C. The samples were immediately incubated for 2 min on ice before they were separated on a MOPS agarose gel containing 1.2 g agarose, 5.5 ml 20% MOPS buffer (400 mM MOPS, 100 mM sodium acetate, 20 mM EDTA) and filled up to 100 ml distilled water. After the separation, the 16 and 23S rRNA loading control were detected using UV-light. To transfer the RNA on a positively charged nylon membrane, vacuum blotting was carried out for 1.5 h and 5 bars. After UV-crosslinking of the membrane, the membrane was prehybridized for 1 h at 42°C in prehybridization buffer containing 20 ml formamide, 10 ml 20x SSC buffer (3 M NaCl and 0.3 M NaCitrate, pH 7), 8 ml 10x blocking agent (Roche Blocking reagent), 2 ml N-Laurylsarcosinate (20 mg/ml), 40 µl 20x SDS, and 2 ml distilled water. 3 µl of the DIG-labelled DNA probe was added to 200 µl water, heated (10 min, 95°C), and cooled down (5 min on ice) twice a row. After a final heating to 95°C for 10 min, the probe was directly added to 20 ml prehybridization buffer. The nylon membrane was incubated and hybridized with the DNA probe overnight at 42°C. Next, the membrane was washed twice for 5 min at room temperature with washing buffer 1 (0.2x SSC buffer, 0.1% SDS) followed by an additional washing step for 15 min at 68°C with washing buffer 2 (0.1x SSC buffer, 0.1% SDS). The membrane was then blocked for 1 h at room temperature in 1x blocking reagent (Roche Blocking reagent) in maleic acid buffer. To visualize the *hcp4* transcript, the membrane was incubated for 1.5 h with a α-Digoxygenin antibody (Anti-Digoxigenin-AP (fab fragments), Roche) 1:6000 in 1x blocking agent in maleic acid buffer. After the incubation with the antibody, the membrane was washed twice for 15 min each at room temperature with CPD* washing buffer (0.1 M maleic acid, 0.15 M NaCl, 0.3% TWEEN-20, pH 7.5). Washing of the membrane was subsequently followed by equilibration for 5 min in the detection buffer (0.1 M Tris-HCl pH 9.5, 0.1 M NaCl). To finally detect the hybridized probe, the membrane was incubated for 5 min with 1 ml of the substrate solution (1:100 CDP* (CDP-Star system, Roche) in detection buffer). The signal was recorded via exposure on X-ray films (CL-Xposure, Thermo Scientific, USA).

### Western blot

For the detection of RovC, Hcp4 and ClpV4-GFP proteins, cultures were incubated depending on the desired conditions. At indicated timepoints, whole cell extracts were prepared by pelleting 1 ml of each sample for 5 min at 8,000x g at RT. The pellet was resuspended in an adjusted 1x Laemmli buffer [126] (40% v/v glycerol, 240 mM Tris-HCl, 8% w/v SDS, 5% v/v β-mercaptoethanol and 0.04% w/v bromophenol blue) to an OD_600_ of 10 and heated for 10 min at 95°C. Proteins were separated on a 15% polyacrylamide gel and transferred onto a polyvinylidene fluoride (PVDF) membrane (Sigma) by Western blotting. Primary antibodies against RovC (1:1000 dilution) and Hcp4 (1:5000) were generated by Davids Biotechnology, ClpV4-GFP was visualized using a monoclonal antibody against GFP (#MAB 3580, Merck Millipore, 1:2000). GAPDH was used as loading control and visualized with a monoclonal antibody against bacterial GAPDH (#MA5-15738, Invitrogen, 1:2000).

### ß-galactosidase assays

The ß-galactosidase assay, first described by Miller 1972 [127], was used to determine *rovC* promoter activity. *Y. pseudotuberculosis* strains harbored pAKH189 or pTS03, respectively were incubated at 25°C and 37°C. At indicated timepoints, 200 µl samples were permeabilized by adding 50 µl 0.1% SDS and chloroform, respectively. After incubation for 10 min, 1.8 ml Z-buffer was added (60 mM Na_2_HPO_4_, 20 mM NaH_2_PO_4_, 10 mM KCl, 1 mM MgSO_4_). The reaction was started by adding 400 µl ONPG as substrate (4 mg/ml). The reaction was stopped by adding 1 ml of 1 M Na_2_CO_3_. The activities were calculated as follows: ß-galactosidase activity [Miller units] = 1000 x OD_450_ x Δt (min)^-1^ x V (ml)^-1^ x OD_600_.

## Acknowledgments

We would like to thank Dr. Ann-Kathrin Heroven for constructing the *lon* deletion mutant. We also thank the members of the Institute of Infectiology for helpful discussions throughout the course of this project.

## Supporting information

**Table S1: Bacterial strains and plasmids**

**Table S2: Oligonucleotides for DNA amplification**

S1 Fig. *clpV4-gfp* expression is repressed at 37°C. Fluorescence microscopy of *Y. pseudotuberculosis* wt *clpV4-gfp* incubated for two and 6 h at 25°C (A) or 37°C (B). Representative images of the brightfield and GFP channels and an overlay of both channels were shown. Bacteria were imaged on agarose pads containing 1% agarose, and the scale bar represents 10 µm.

S2 Fig. Gating strategy to analyze heterogeneous T6SS4 and *rovC* expression. (A) Samples were stained with DAPI to gate for bacteria. (B) PI staining to exclude dead bacteria. (C) Negative control (YPIII wildtype) for GFP expressing bacteria. (D) Gate for GFP-expressing bacteria. (E) Negative control (YPIII wildtype) for mCherry expressing bacteria. (F) Gates for mCherry-expressing bacteria, divided into low and high expression intensity. H=height, A=area, SSC=side scatter. Exemplary plots are shown.

S3 Fig. Deletion of *lon* results in increased *rovC* promoter activity. Wt *clpV4-gfp* p*P_rovC_ rovC’-‘mCherry* and Δ*lon clpV4-gfp* p p*P_rovC_ rovC’-‘mCherry* were incubated overnight at 25°C. 1 x 10^5^ bacteria were analyzed with flow cytometry, and experiments were performed in four independent experiments. Statistical significance was tested with an unpaired t-test. ns = not significant p > 0.05.

S4 Fig: T6SS4 gene expression is downregulated at 37°C. (A-C) Total RNA of an overnight culture of wt, Δ*csrA,* and Δ*ymoA* was isolated to perform qRT-PCR. Specific primer pairs for five T6SS4 genes were used to determine the expression within the T6SS4 operon. Log2-fold changes were calculated between the T6SS4 transcript and *sopB* as a non-temperature-regulated reference gene [6]. Experiments were performed in three biological replicates, and significant differences were determined using a Two-Way ANOVA with Šidák correction **** = p ≤ 0.0001

S5 Fig. The *rovC* promoter activity is not subject to temperature-dependent control. Fluorescence microscopy of *Y. pseudotuberculosis* wt *clpV4-gfp* harboring *pP_rovC_rovC’-‘mCherry*. Strains were incubated for 2 h and 6 h at either 25°C (A) or 37°C (B). Representative images of the GFP and mCherry channels, overlays of the GFP and mCherry channels, and an overlay of all channels with the brightfield, were shown. Bacteria were imaged on agarose pads containing 1% agarose, and the scale bar represents 10 µm.

S6 Fig. Accumulation of *hcp4* transcript is not due to an additional *hcp4* promoter. A low copy plasmid harboring the upstream region of *hcp4*, p*hcp4’-‘gfp* (-212 to +30 base pairs with respect to the translational start site) was transformed into YPIII wt. Samples were incubated in 20 ml LB overnight at 25°C. wt *clpV4-gfp* was used as a control to compare the amount of *clpV4-gfp* expressing bacteria to the potential *hcp4* promoter activity. Samples were taken for flow cytometry, and 1 x 10^5^ cells were analyzed. Experiments were performed in three biological replicates. The data depict the mean and standard deviation. pV = empty vector control.

S7 Fig. *hcp4* transcript is the most abundant within the operon. RNA coverage of all T6SS4 genes from samples of YPIII wt, incubated for 2 h at 25°C. Data was taken from Meyer *et al*., 2024 [46].

